# Characterization of a cofilin mutant with high actin bundling activity in living cells

**DOI:** 10.64898/2026.04.22.720186

**Authors:** Bruno F. Pizani, Leanna M. Dover, Michelle Cobb, Jarrett Lloyd, Jordan M. Hardeman, Karen A. Litwa, Robert M. Hughes

## Abstract

Cofilin is a key regulator of actin dynamics that, along with a myriad of other actin-binding proteins, controls the balance of F- and G-actin in numerous cell types. While prior structural studies of the cofilin-actin binding interface have delineated many critical interactions between cofilin and actin, the roles of some residues within the cofilin-actin binding interface remain poorly defined. In this study, we investigate the role of cofilin S119 in the cofilin-actin interaction. Despite its unique position within the cofilin-actin interface and its putative role as a phosphorylation site, relatively little direct evidence exists to define whether it plays an important role in cofilin-actin dynamics. Using site-directed mutagenesis, we demonstrate that mutation of S119 to aromatic amino acids (W, F, Y) results in cofilins with strong actin bundling activity in living cells. This activity can be countered by the incorporation of mutants that disfavor actin rod forming activity (R21Q). Mutation of S119 to phospho-mimic (E) and non-phosphorylated (A) residues either strongly inhibits (E) or modestly increases (A) actin bundling activity. Expression of the S119W mutant in neurons reveals its impacts on spine length and size, while FRAP studies show that its mobile fraction is intermediate between that of LifeAct and WT cofilin. Finally, it is shown that the strong actin bundling phenotype associated with S119W inhibits the progression of optogenetically induced apoptosis.

## INTRODUCTION

The actin binding activity of cofilin is regulated by several key residues, including Ser 3, Ser 94, and Lys 96 (1–3). Mutagenesis of these residues can greatly reduce or eliminate cofilin’s actin binding and severing activity (3). More recently, molecular modeling studies of the cofilin-actin binding interface have elucidated the position of these residues within the cofilin-actin interface while identifying additional residues that may also be critical for actin binding (2). One such residue, S119, was found to be uniquely positioned within the cofilin-actin interface (**Fig. 1**), where it points directly into the actin interface and having close contacts with actin residues Y143 and L346 (2). Beyond its prior identification in a bioinformatics survey as a putative phosphorylation site (4, 5), and as an important residue in the interaction of cofilin and LIM kinase (6), its role in actin binding remains less clear. In this study, a site-directed mutagenesis approach has been used to demonstrate that mutation of Ser 119 to aromatic amino acids results in cofilin mutants with strong actin-bundling phenotypes (in particular S119W), producing structures that closely resemble cofilin-actin rods. A mutagenesis approach has also shown that the actin bundling activity of Ser 119 mutants can be countered with the addition of the cofilin-actin rod resisting mutant R21Q (7), and that phosphorylation-mimicking mutation S119E eliminates actin bundling. The functional properties of the S119W mutant are further demonstrated through FRAP analysis of actin binding dynamics and through comparative studies of apoptotic progression using an optogenetic tool. Overall, these results point to an important regulatory role of Ser 119 in cofilin-actin binding that adds to the list of key residues within the actin-cofilin interface while inviting further inquiry into its role in regulating actin dynamics and cellular behavior.

**Figure 1.**
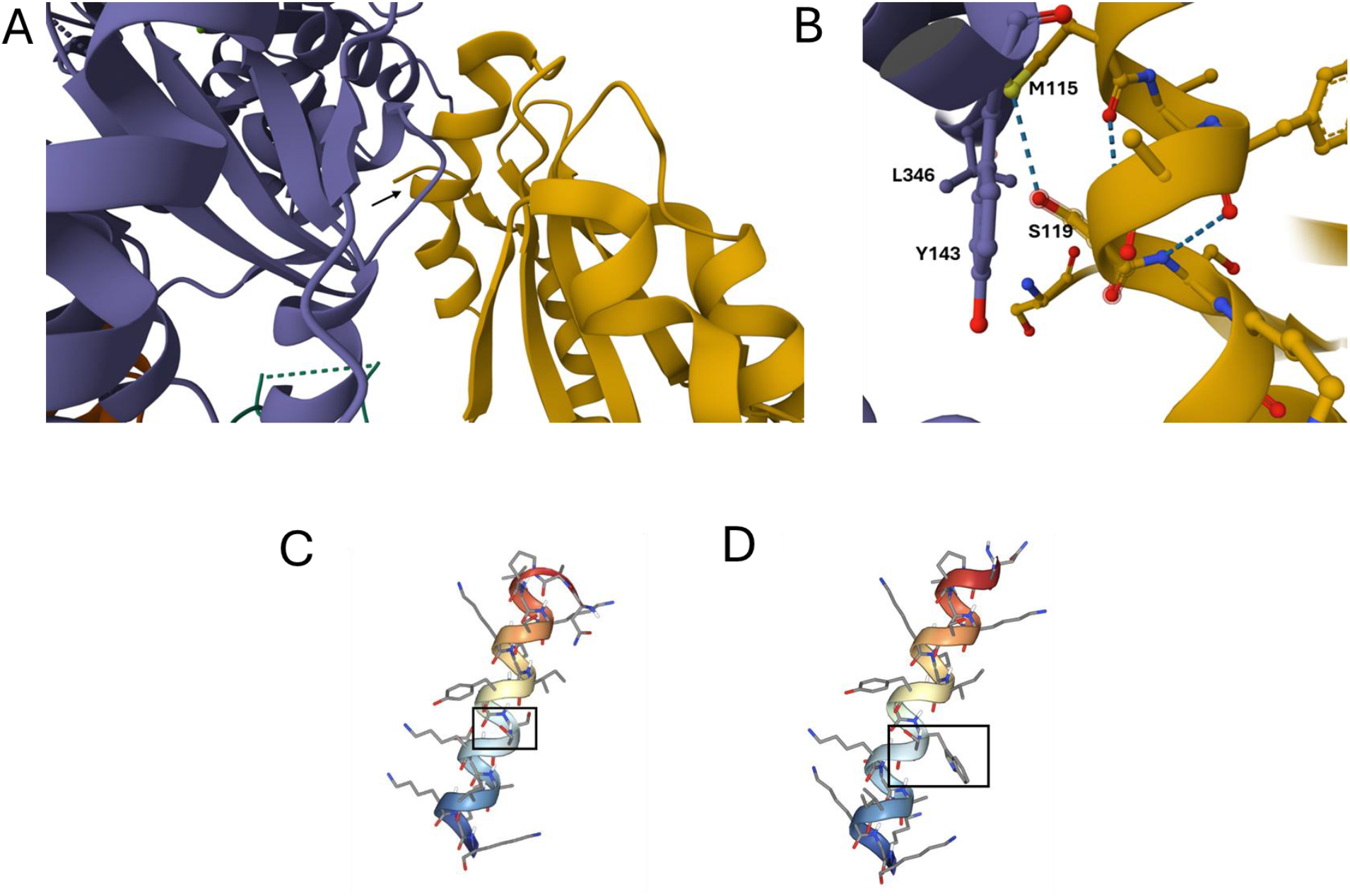
Structure of cofilin-actin binding interface reveals a potentially unique role for cofilin S119. A. Actin (purple) and cofilin (gold) interface from structure of cofilin-decorated actin filament (RCSB PDBID 5YU8 (1)). Black arrow points to location of cofilin S119. B. Zoomed-in region of actin-cofilin binding interface. Cofilin S119 has a hydrogen bond to cofilin M115 and close contacts with actin Y143 and L346. C. PEP-FOLD (26)model of cofilin helix with S119 residue outlined with black rectangle. D. PEP-FOLD (26)model of cofilin helix with S119W mutation.

## RESULTS AND DISCUSSION

Previous molecular modeling-based studies of the cofilin-actin binding interface indicated that cofilin residue Ser119 is uniquely positioned within the cofilin-actin interface to be a potential driver of actin-cofilin interactions (2). In these modelling studies, the cofilin residue Ser 119 located within alpha helix 4 of cofilin had 100% contact with the actin filament throughout simulations of actin-cofilin binding (2). By contrast, the adjacent Ser 120 residue only contacted the actin filament 13% of the time (2), indicating that Ser119 may play a previously underappreciated role in regulating actin dynamics. Given these results, along with the existing structural data on cofilin bound to F-actin filaments (**Fig. 1A**) and the predicted structural integrity of cofilin helix 4 (**Fig. 1B**), we considered whether the mutation of the Ser 119 residue might significantly alter cofilin-actin binding. To test this, we mutated Ser 199 to Trp, in a cofilin-GFP fusion protein, anticipating that this hydrophobic, planar amino acid might pack well into this unique location within the cofilin-actin interface. Transfection of the resulting mutant (cofilin S119W) in HeLa cells produced a remarkable actin-bundling (cofilin-actin rod) phenotype (**Fig. 2**), which is readily controlled by varying the quantity of transiently transfected DNA. We note that it has long been observed that even transient overexpression of wild type cofilin can result in a high level of cofilin-actin rod formation (7, 8). Nonetheless, the number of cofilin-actin rods was greater with the S119W mutant versus WT cofilin at all DNA concentrations tested with a maximum difference apparent at the 250 ng transfection level (**Fig. 2A-B**). We confirmed that the observed structures consisted of both cofilin and actin using a dual transfection experiment with mCherry-actin (**Fig. 2C**).

**Figure 2.**
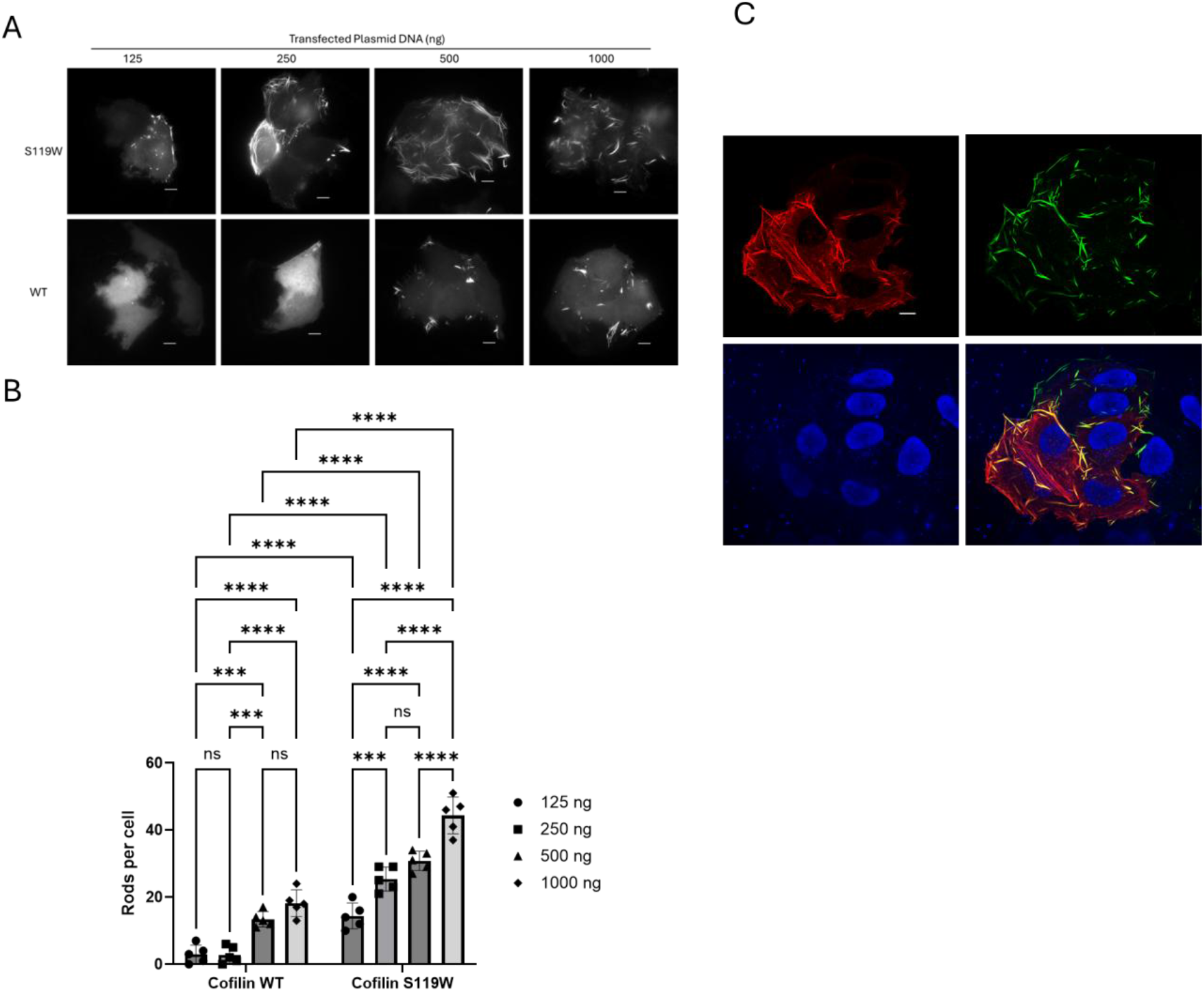
Mutation of cofilin serine 119 to tryptophan (S119W) promotes a cofilin-actin rod phenotype in HeLa cells. a. Widefield fluorescence microscope images of HeLa cells transfected with varying amounts of plasmid DNA encoding Cofilin.S119W.GFP or Cofilin.WT.GFP. Scale bars = 10 microns. B. Quantification of rods/cell for each titration condition. n=5 cells assessed for rod content for each experimental condition (TwoWay ANOVA: ***, p = 0.0001; ****, p< 0.0001, ns = not significant). C. Confocal fluorescent microscope images of Hela cells transfected with Cofilin.S119W.GFP and mCherry-actin. Images show mCherry-actin (upper left panel), Cofilin.S119W.GFP (upper right panel), Hoescht 33342 (lower left panel), and overlay (lower right panel). Scale bars = 10 microns.

To further characterize this mutation site, we investigated whether other aromatic amino acids might have the same effect as tryptophan (**Fig. 3**) and how other phosphorylation-state mimicking mutations (Ser −> Ala for non-phosphorylation and Ser −> Glu for phosphorylation) would impact the observed cofilin-actin phenotype. Both aromatic amino acids (Phe, Tyr) investigated had rod forming properties similar to those of S119W, and all exhibited rod formation greater than that of the S119A non-phosphorylated actin mutant (**Fig. 3A-B**). By contrast, the phospho-mimic S119E mutant exhibited no cofilin-actin rod formation (**Fig. 3A-B**). The S119A/S119E mutant data indicates that S119 may play a role in the control of cofilin activity. While S119 has been indicated as a putative phosphorylation site in bioinformatics surveys of cofilin function at the single residue level (4) it is not frequently described in the literature as such. Notably, prior work has identified S119 a mediator of cofilin-LIM Kinase binding (6), indicating that it could be part of the regulatory network between LIMK regulation of actin binding. As a result, further inquiry into the role of S119 as a node in the control of cofilin-actin dynamics is warranted.

**Figure 3.**
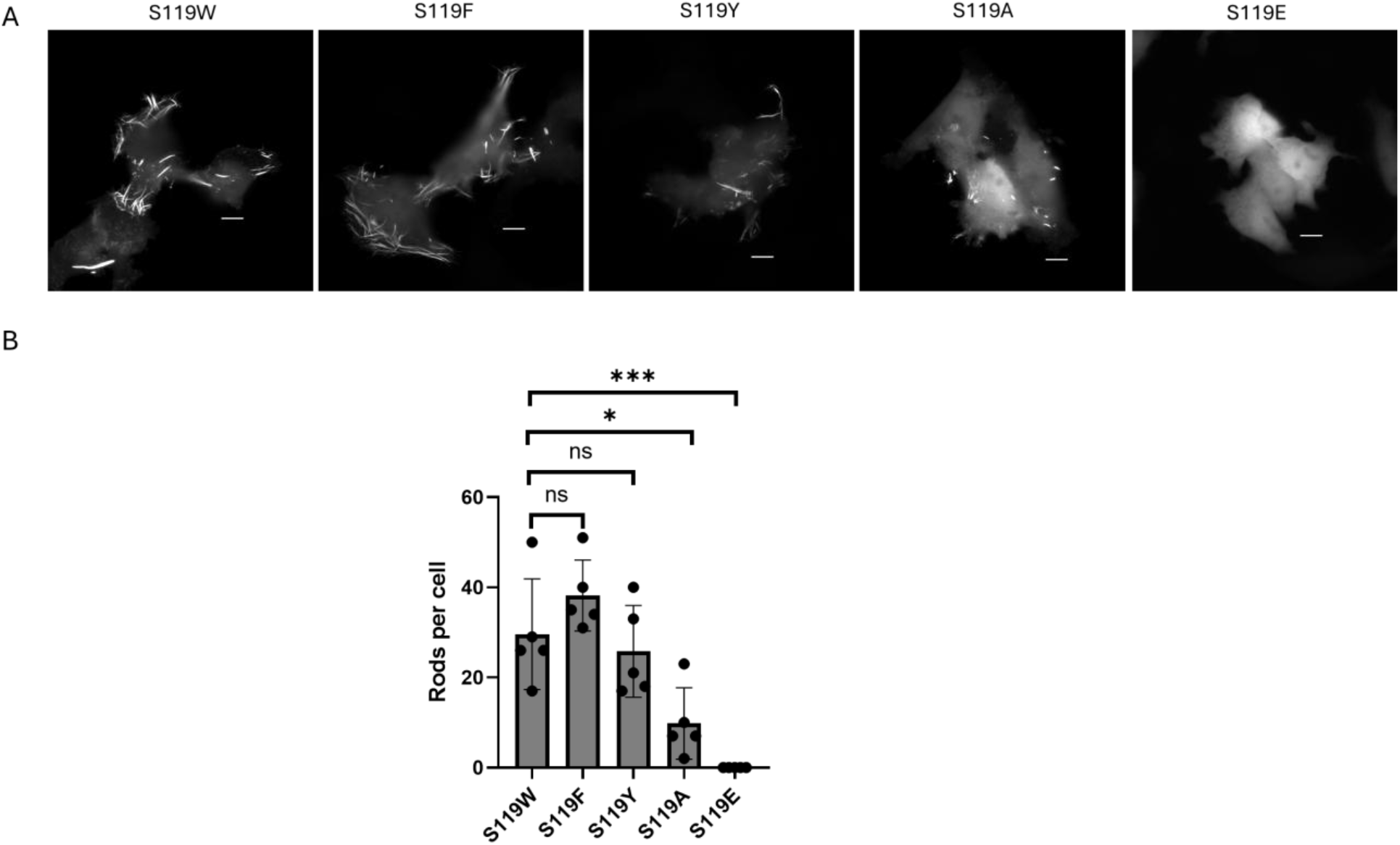
Mutation of cofilin Ser 119 to aromatic amino acids and phosphorylation state mimicking amino acids. a. Widefield fluorescence microscope images of HeLa cells transfected with various Ser 119 mutants of Cofilin.GFP (250 ng per transfection). Scale bars = 10 microns. B. Quantification of rods/cell for each titration condition. n=5 cells assessed for rod content for each experimental condition (OneWay ANOVA: *, p = 0.0137; ***, p=0.0003; ns = not significant).

Cofilin-actin rod formation is of interest for its possible role in neurodegenerative disease progression (8–14). To aid in the investigation of cofilin’s role in rod formation, Bamburg and colleagues created a novel cofilin mutation designed to resist incorporation into cofilin-actin rods in the absence of applied cellular stress (7). This mutant (cofilin R21Q), expressed as a fusion to GFP, was shown to have no propensity for cofilin-actin rod formation in living cells under conditions of homeostasis (7). To investigate whether this mutation would impact the actin bundling observed with S119W, we created a cofilin R21Q-S119W double mutant and R21Q single mutant for comparison in our expression system. As expected, the R21Q single mutant demonstrated negligible rod formation, while the R21Q-S119W double mutant exhibited greatly reduced rod formation in comparison to the S119W mutant (**Fig. 4**). By contrast, introducing the constitutively active S3A mutant alongside the S119W mutation had relatively little impact on the abundance of cofilin-actin rods. The ability of the R21Q mutation to mitigate the impacts of S119W points to a hierarchy of cofilin-actin filament interactions, arranged not only by location (R21 is located on an unstructured loop of cofilin and interacts with a different region of actin) but by interaction type (R21Q eliminates an electrostatic interaction with the actin interface whereas S119W promotes filling of a hydrophobic pocket).

**Figure 4.**
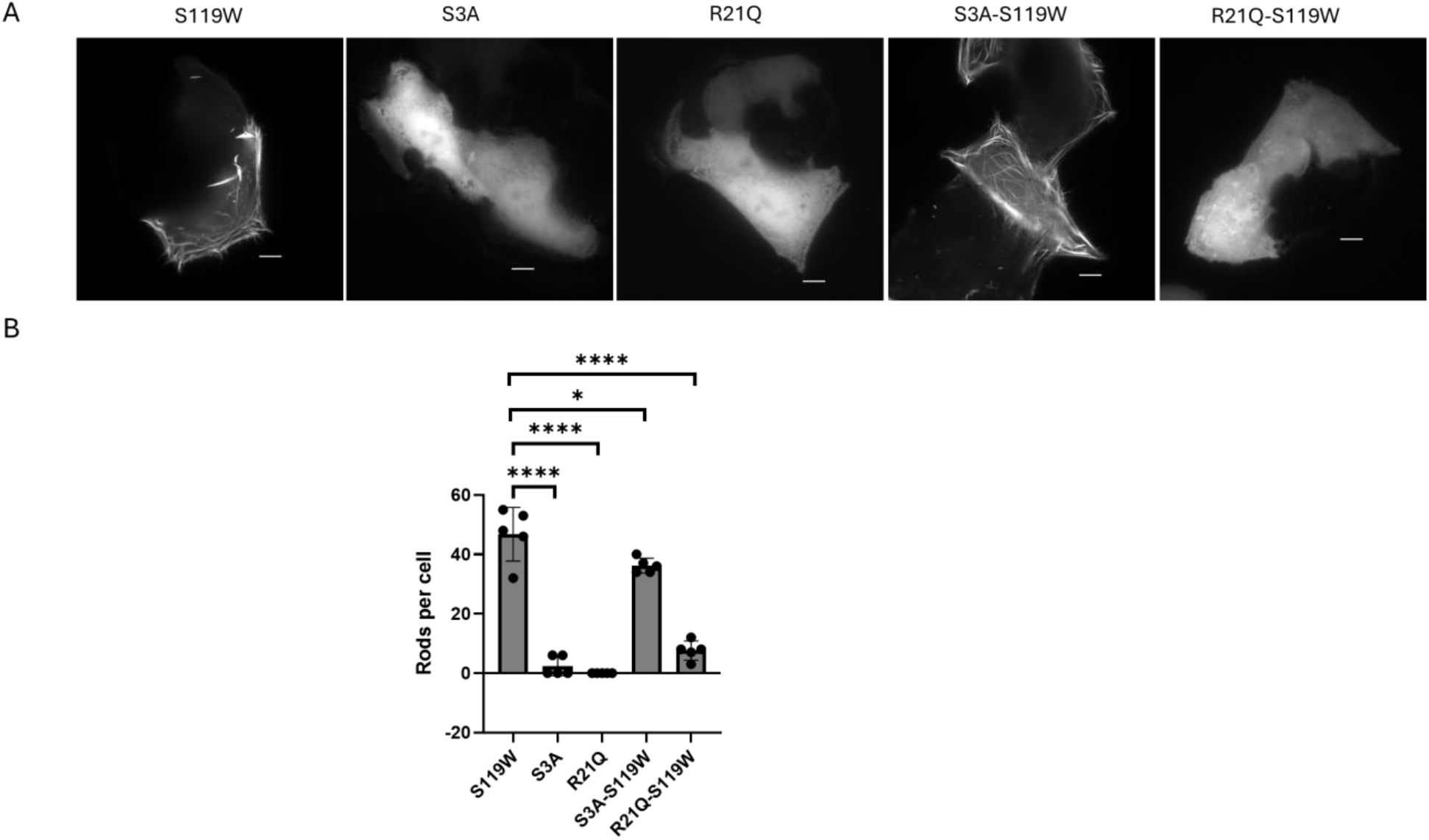
Rod-resistant mutation R21Q reverses actin bundling phenotype of S119W. a. Widefield fluorescence microscope images of HeLa cells transfected with various mutants of Cofilin.GFP (250 ng per transfection). Scale bars = 10 microns. B. Quantification of rods/cell for each titration condition. n=5 cells assessed for rod content for each experimental condition condition (OneWay ANOVA: *, p = 0.140; ****, p< 0.0001).

Given the association of cofilin-actin rod formation with neurodegeneration, we further evaluated the role of cofilin S119 in neuronal physiology (**Fig. 5**). Dendritic spines on neurons are actin-enriched structures, whose dynamics depend on actin-regulation. In response to NMDA receptor activation, spines undergo morphological changes associated with long-term potentiation that facilitate action potential formation, notably stably increasing the volume of the spine head. During the initial phase of LTP, cofilin is initially recruited in its non-phosphorylated active form to facilitate actin-remodeling associated with spine head expansion (15, 16). However, cofilin phosphorylation is necessary for subsequent stabilization of the increased spine volume, underlying the consolidation of learning and memory events (15, 16). Not surprisingly, cofilin-actin rod formation inhibits synaptic plasticity necessary for LTP, driving memory loss and neurodegeneration. To evaluate cofilin S119’s role in spine dynamics, we transfected primary mouse neurons with WT-cofilin, cofilin-S119W, S119A, S119E and Lifeact, the original genetically-encoded cofilin-based sensor used to evaluate spine-associated actin dynamics (17, 18). Consistent with the observed actin bundling phenotype in HeLa cells, cofilin S119W exhibited numerous long filopodia-like spikes emanating from neurites, similar to expression of WT-cofilin, Lifeact, and cofilin-S119A (**Fig. 5A-B**). Notably, expression of cofilin-S119E resulted in spines that most closely resembled those observed in the GFP control group, with many beginning to mature and exhibit a spine head. To compare how the cofilin S119W mutant altered cofilin dynamics, we use fluorescence recovery after photobleaching (FRAP) to assess cofilin mobility (**Fig. 5**). While all cofilin constructs evaluated exhibited a similar half-life of recovery, cofilin S119W and cofilin s119A significantly decreased spine-associated cofilin motility when compared to Lifeact, albeit to a lesser extent than expression of WT cofilin (**Fig. 5C**). This reduced motility suggests that mutation of S119 to either the aromatic residue tryptophan or to alanine to mimic the non-phosphorylated state increases actin association. Conversely, the phospho-mimetic cofilin S119E, exhibited motility similar to Lifeact, suggesting decreased actin association relative to S119W. All constructs exhibited similar rates of recovery. Together, S119W’s increased formation of spiky dendritic spines and decreased motility, suggest that cofilin S119W similarly regulates actin association in neurons as it does in HeLa cells, and that S119 phosphorylation could play a role in regulating actin association similar to the canonical phosphorylation of cofilin at Ser3 (3).

**Figure 5.**
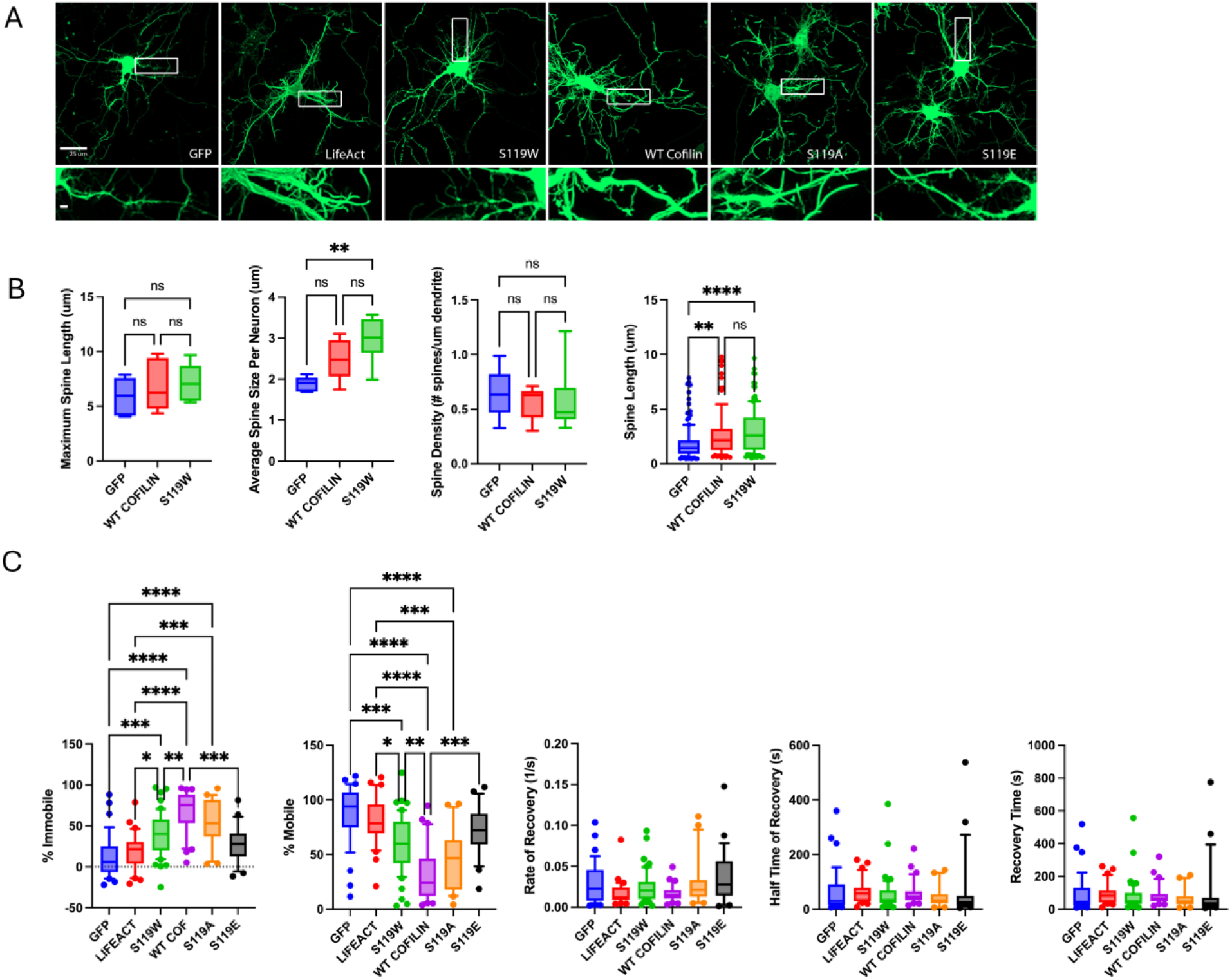
FRAP of S119W mutant in neurons reveals stronger actin binding activity than Lifeact. A. DIV-8 mouse primary cortical neurons expressing GFP-labeled cofilin constructs prior to photobleaching were used to assess changes in immature spine length and density. **B**. Both WT cofilin and cofilin S119W significantly increased spine length, but not spine density. However, only cofilin S119W significantly increased spine length when spine length was averaged across experiments. **C**. To analyze the kinetics of the various cofilin constructs, we performed fluorescence recovery after photobleaching in dendritic spines ~1 week old mouse primary cortical neurons (between DIV7-DIV11). Cofilin-S119W, WT-cofilin and cofilin S119A all significantly increased the immobile fraction and resulted in a corresponding decrease in the mobile fraction, when compared to either GFP or Lifeact, suggesting increased actin association. However, cofilin-S119W, was also significantly different from WT-cofilin, exhibiting mobility intermediate between Lifeact and WT cofilin which had the greatest immobility. The phosphomimetic cofilin-S3119E exhibited kinetics similar to GFP and Lifeact, indicative of decreased actin association.

Finally, we investigated the functional consequences of enhanced actin bundling with the cofilin S119W mutation using an optogenetic activator of apoptosis (OptoBax) (19–21). In previous studies, we have demonstrated that pharmacological inhibition of F-actin turnover inhibits apoptotic progression (19). Furthermore, cofilin-actin rod formation occurs during apoptosis and may comprise a cellular defense mechanism that preserves energy by halting actin dynamics (22, 23). In this experiment, we investigated whether cells expressing S119W and exhibiting a strong cofilin-actin rod phenotype affected the onset of OptoBax-mediated apoptosis (**Fig. 6**). In these experiments, HeLa cells exhibiting the actin bundling phenotype were better able to resist apoptosis over a 2 hour period while cells expressing the S119W mutant but not exhibiting a strong cofilin-actin rod phenotype generally entered apoptosis-associated cellular collapse. This demonstrates that perturbation of actin dynamics via overexpression of the cofilin S119W mutant can have profound impacts on cell signaling pathways.

**Figure 6.**
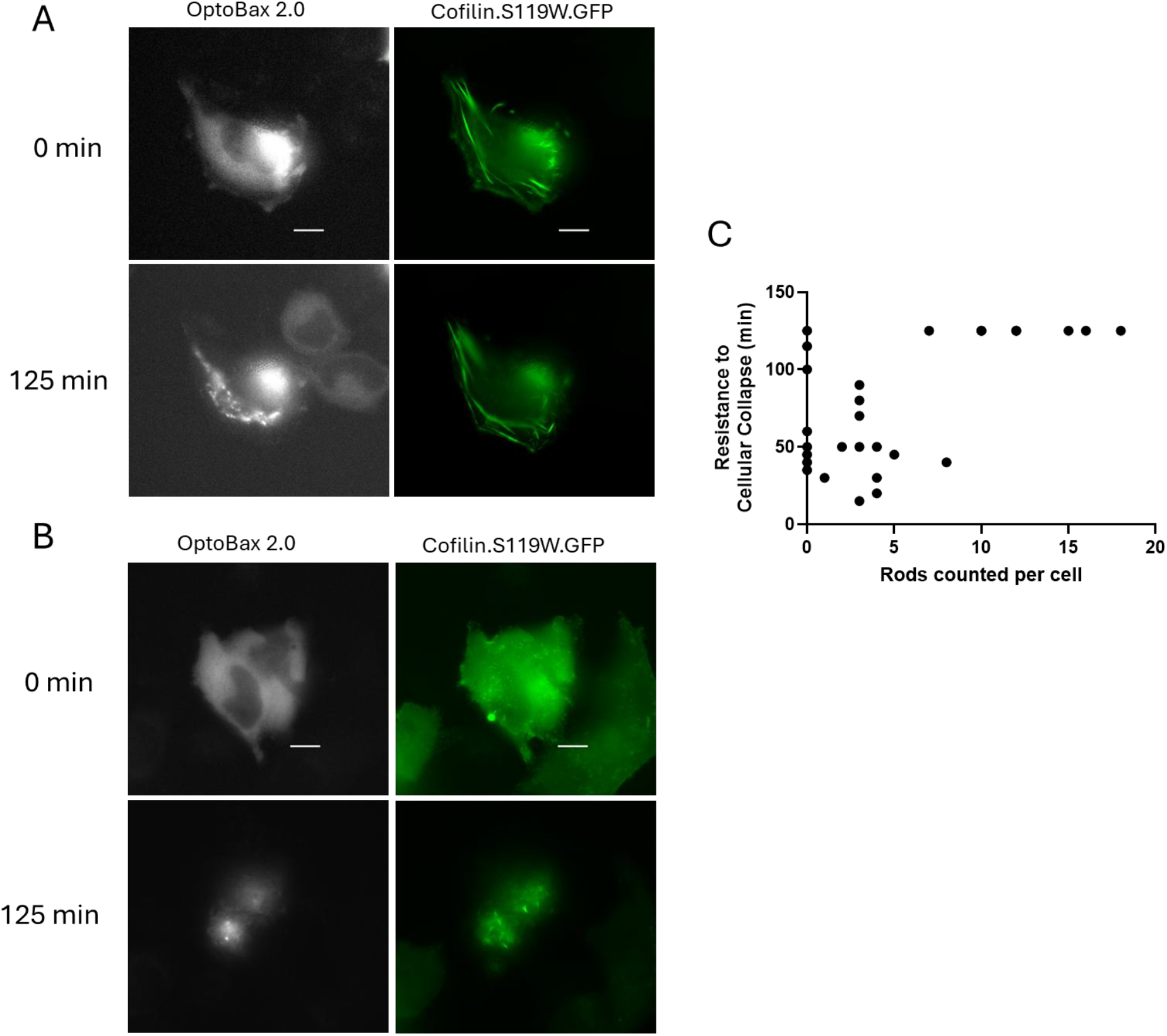
Presence of cofilin S119W mutant-induced actin bundles inhibits progression of apoptosis in an optogenetic apoptosis system. A. HeLa cells co-transfected with OptoBax 2.0 and Cofilin.S119W.GFP with increased cofilin actin rods. Scale bars = 10 microns. B. HeLa cells co-transfected with OptoBax 2.0 and Cofilin.S119W.GFP with no mature rods. Scale bars = 10 microns. C. Plot of number of rods per cell versus time to apoptosis-associated cellular collapse. All cells with 10 or more rods per cells resisted collapse for the duration of the experiment (125 minutes).

## CONCLUSIONS

In this work, we investigated whether the mutagenesis of Cofilin Ser 119 impacts cofilin-actin binding and actin dynamics. Mutation of cofilin Ser 119 to tryptophan created a strong actin bundling (rod) phenotype in cells while phenotypes resulting from mutation to phosphorylation-state mimics indicate that Ser 119 could be an important node for cofilin control via phosphorylation. These results suggest that additional confirmation of Ser 119 as a functional node for cofilin control are warranted as limited studies exist in the literature. Intriguingly, cofilin Ser 119 has been implicated in binding of LIM kinase, a critical kinase for control of cofilin activity through phosphorylation at Ser 3, and thus Ser 119 might be important for multiple protein-protein interactions involving cofilin. This study corroborates the identification of Ser 119 within the cofilin-actin binding interface as a residue uniquely positioned to interact with actin consistently throughout dynamic cofilin-actin binding and demonstrates, through inhibition of apoptosis, that in vivo changes in this residue either through post-translation modification or genetic mutation could have implications for cell signaling and neurodegenerative disease progression. We propose that the inclusion of cofilin Ser 119 as a control in studies of cofilin-actin rod formation and dynamics might be beneficial to the field as current methods for the induction of cofilin-actin rods rely upon the application of cellular stressors, including inhibitors of glycolysis and electron transport. This mutant presents a means of introducing abundant rods into cells without requiring exogenous perturbation of the cell’s energy production system (24, 25).

## Funding and additional information

This work was supported by a grant from the National Institutes of Health (grant no.: 1R15NS125564-01) to R.M.H. and a grant from the National Science Foundation through CAREER award number 2144912 to K.A.L. The content is solely the responsibility of the authors and does not necessarily represent the official views of the National Institutes of Health or the National Science Foundation.

## MATERIALS AND METHODS

### Plasmids and Cloning

Cloning of the Cofilin.GFP construct conducted according to previously reported methods (27, 28). Briefly, a gene encoding Cofilin 1 (M. musculus) was PCR amplified with primers appending XhoI and HindIII restriction sites. Amplified gene fragments were trimmed via restriction digest, followed by ligation into complementary restriction sites in the target plasmid (phCMV-GFP (Genlantis)). Point mutations were introduced into the plasmid encoding Cofilin-GFP using mutagenic primers (IDT DNA) following a standard protocol for site-directed mutagenesis. mCherry-Actin-C-18 was a gift from Michael Davidson (Addgene plasmid # 54967) (29). Cloning and validation of OptoBax 2.0 constructs has been described elsewhere (19).

### Cell lines and transfection

Transfection of HeLa cells was then performed with Calfectin reagent (SignaGen) following the manufacturer’s suggested protocols. Briefly, for single transfections in 35 mm glass bottom dishes, plasmid DNA was added to 100 µl of DMEM, followed by the addition of 3 µl of Calfectin reagent. For dual transfections (Cofilin-GFP with mCherry-Actin), plasmid DNA (1000 ng per construct) was combined in a 1:1 ratio. For triple transfections (Cofilin-GFP with OptoBax 2.0 (Cry2(1-531).mCherry.BaxS184E and Tom20.CIB.GFP), plasmid DNA was combined in a 1:1:0.25 ratio (1000 ng Cry2.mCherry.Bax.S184E; 1000 ng Tom20.CIB.GFP; 250 ng Cofilin.S119W.GFP). The solution was incubated at room temperature for 10 min, followed by dropwise addition to cell culture. Transfection solutions were allowed to remain on cells overnight. Cells were maintained at 37°C and 5% CO2 in a humidified tissue culture incubator in a culture medium of DMEM supplemented with 10% FBS and 1% Penicillin-Streptomycin. Cell media was replaced with L-15 containing 10% FBS immediately prior to imaging.

### Neuron cultures and transfection

Primary neuron cultures were performed as previously described (30). Cortices were dissected from embryonic day (E)18.5 mouse brains and were dissociated following an protocol adapted from Bunner et al. (24) using a primary neuron isolation kit (Thermo Scientific™ Neuron Isolation Enzyme (with papain) for Pierce™ Primary Cell Isolation Kits Catalog No. PI88285) and triturated in growth media with serum and then transferred to a solution of warm Opti-MEM | Reduced Serum Medium, with glucose (Gibco/Fisher Scientific, Cat# 31-985-062) for centrifugation. Isolated neurons were resuspended in fresh growth media with serum and plated onto 12-well glass bottom dishes for imaging. After 2 days in vitro (DIV), the cultures were changed to Prenatal Mouse Neuron Media containing Neurobasal media (Gibco/Fisher Scientific, Cat# 21-103-049) supplemented with B-27 Plus (Fisher Scientific, Cat# A3582801), GlutaMAX Supplement, and penicillin/streptomycin. Half of the media was changed every other day. Neurons were transfected between div 4-8 with 1ug DNA using either lipofectamine 2000 of lipofectamine LTX (Thermo Fisher Scientific).

### FRAP experiments and quantification

Live imaging was performed on a Zeiss LSM 800 microscope with continuous recording for 5 min. Fluorescence photobleaching was performed using 100% 488nm laser power and repeated bleaching to reduce fluorescence intensity in the selected ROI by 10% of the initial intensity. Live imaging was acquired using definite focus to minimize z fluctuations. Following image acquisition, we used the Zen Blue software to process and analyze the resulting timelapse recordings, by first performing timelapse alignment to minimize x,y drifts, followed by use of the FRAP analysis module to analyze the resulting kinetics. A reference unbleached area was selected as was a background region. The software fitted the resulting data to the following equation, I = I_E_ – I_1_*exp(-t/T1), where I is the fluorescent intensity at a given time, I_E_ is the ending fluorescent intensity, I_1_ the amplitude of intensity recovery, t is the given time from the photobleaching event and T1 is the time constant. We excluded resulting data that was less than −20% mobility or greater than 120% mobility as these errors often resulted from spine retraction or formation within the region of interest that interfered with the detection of fluorescence recovery.

### Confocal Microscopy

Confocal images of fixed cells were obtained with a Zeiss LSM 800 microscope with Airyscan technology. Fluorescence images were colorized and overlaid using FIJI software.

### Widefield Microscopy

A Leica DMi8 Live Cell Imaging System, equipped with an OKOLab stage-top live cell incubation system, LASX software, Leica HCX PL APO 63x/1.40-0.60na oil objective, Lumencor LED light engine, CTRadvanced+ power supply, and a Leica DFC900 GT camera, was used to acquire images. Exposure times were set at 100 ms (GFP, 470 nm) with the LED light source at 50% power. For time-resolved optogenetic experiments, exposure times were set at 50 ms (GFP, 470 nm) and 200 ms (mCherry, 550 nm), with LED light sources at 50% power, and images acquired every 5 minutes over a 125-minute time course.

### Rod counting

Analysis of imaging data was performed in FIJI, equipped with the BioFormats package (31). Rods were counted by choosing one representative cell from each field of view. Rods were selected based on linear appearance (long, unobstructed filaments). For regions with branched rods, each branch was counted as an individual rod. Small, circular aggregates were not included in the analysis.

### Statistical analyses

Statistical significance (p values) were determined using GraphPAD Prism (Prism 10 for Windows Version 10.6.1 (892)). One-way and Two-way ANOVAs were used to analyze rod counts in HeLa cells (32). Plots were constructed using GraphPAD Prism version (10.6.1).

